# A kinetic model for USP14 regulated substrate degradation in 26S proteasome

**DOI:** 10.1101/2024.12.31.630858

**Authors:** Di Wu, Qi Ouyang, Hongli Wang, Youdong Mao

**Author notes:** Corresponding author (HW), (YM).

## Abstract

Despite high-resolution structural studies on the USP14-proteasome-substrate complexes, time-resolved cryo-electron microscopy (cryo-EM) results on USP14-regulated allostery of the 26S proteasome are still very limited and a quantitative understanding of substrate degradation dynamics remains elusive. In this study, we propose a mean field model of ordinary differential equations (ODEs) for USP14 regulated substrate degradation in 26S proteasome. The kinetic model incorporates recent cryo-EM findings on the allostery of 26S proteasome and generates results in good agreement with time-resolved experimental observations. The model elucidates that USP14 typically reduces the substrate degradation rate and reveals the functional dependence of this rate on the concentrations of substrate and adenosine triphosphate (ATP). The half-maximal effective concentration (EC50) of the substrate for different ATP concentrations is predicted. When multiple substrates are present, the model suggests that substrates that are easier to insert into the OB-ring and disengage from the proteasome, or less likely to undergo deubiquitination would be more favored to be degraded by the USP14-bound proteasome.

**Author Summary:** The proteasome is a crucial protein complex involved in the degradation of damaged or unnecessary proteins within cells, requiring ATP and ubiquitin for its functioning. It is regulated by cellular factors that transiently associate with it, often referred to as proteasome-associated proteins. USP14 is such a protein that activates its deubiquitination activity through reversible binding to the proteasome, thereby decreasing the substrate degradation activity of the proteasome. In this study, we developed a kinetic model to describe how USP14 regulates substrate degradation in proteasome based on recent experimental findings. The model yields result consistent with experimental observations, demonstrating that the mean-field description of mass action law for chemical reactions also applies to complex biomolecular machineries.

## 1. Introduction

The proteasome is the core of the ubiquitin-proteasome system (UPS) in eukaryotes, playing a pivotal role in regulating protein degradation processes [1,2]. It is a 2.5-megadalton protein complex composed of core particle (CP) and regulatory particle (RP) [3]. Substrate proteins tagged with ubiquitin are recognized by ubiquitin recognition sites on RPN1 subunit, deubiquitinated by the subunit and unfolded by ATPase motor, translocated into the CP, and ultimately degraded into short peptides at the hydrolytic sites within CP. Previous studies have elucidated the dynamics of the proteasome through various experimental approaches, including mutagenesis experiments [4,5], single-molecule experiments [6,7], and cryo-electron microscopy (cryo-EM) experiments [3,8–10]. In vivo, the function of the proteasome is regulated by cellular factors that transiently associate with it, often referred to as proteasome-associated proteins [11–13]. These include the ubiquitin-specific protease 14 (USP14) [14], deubiquitinating enzyme UCH37 [15–17], E3 ubiquitin ligase UBE3C/Hul5 [18,19], parkin [20], UBE3A/E6AP [21,22], etc. The enzymes interact directly or indirectly with the proteasome and thereby regulate its function. Among these enzymes, the structure and function of USP14 have been the focus of previous studies [14,19,23– 27].

USP14 (or its homolog UBP6 in yeast) is a deubiquitinating enzyme that activates its deubiquitination activity by reversible binding to the proteasome, thereby regulating the UPS system [14]. Biochemical experiments have shown that USP14 can decrease substrate degradation activity of the proteasome [23,25]. Deletion of UBP6 in yeasts can leads to growth deficiencies [19]. Cryo-EM experiments have further studied how USP14/UBP6 binds to the proteasome and modulates its function [24,26,27]. USP14/UBP6 binds to the T2 site on the proteasomal regulatory particle non-ATPase (RPN) 1 via its ubiquitin-like (UBL) domain [28] and cleaves the ubiquitin chains through its ubiquitin-specific protease (USP) domain [25]. Recent time-resolved cryo-EM experiments have identified 13 distinct high-resolution structures of UPS14-bound proteasome and their temporal changes, providing a rough outline of transitions between these complexes [27]. While cryo-EM experiments with temporal resolution on USP14-regulated allostery of the 26S proteasome are very limited and remain a significant challenge, an accurate and quantitative understanding of substrate degradation dynamics is elusive. So far, no modeling studies have been reported regarding the dynamics of proteasomal substrate degradation regulated by USP14.

In this paper, we propose a kinetic model of ordinary differential equations (ODEs) based on recent experimental findings on the regulatory interactions between USP14 and the 26S proteasome during substrate degradation. The model well explains the recent experimental observations on the temporal changes in the distribution of human 26S proteasomal conformations [27]. In consistent with experimental observations, the model illustrates that the proteasome decreases its substrate degradation rate upon the regulation of USP14. Subsequent analyses with simplified models predict how the rate depends functionally on the concentrations of substrate and adenosine triphosphate (ATP), and the substrate’s half-maximal effective concentration (EC50) across a wide range of ATP concentrations. In our model for multiple substrates, USP14 alters the proteasome’s selectivity towards different substrates. Model results predict that the USP14-bound proteasome preferentially degrades substrates that readily access the OB-ring, are easier to detach from the proteasome, and are less prone to deubiquitination. The theoretical model results should aid in achieving a deeper insight into the intricate process of substrate degradation in 26S proteasome regulated by USP14. This is particularly valuable, given the current experimental challenges in obtaining detailed,time-resolved cryo-EM studies of the proteasomal allostery.

## 2. Kinetic model of USP14-regulated proteasomal degradation of substrate

### 2.1 Experimental findings on the human 26S proteasomal conformations

In eukaryotes, the 26S proteasome serves as a central player in the complex process of degradation of proteins. USP14, a ubiquitin-specific protease acting as a deubiquitinating enzyme, binds to the proteasome in a reversible manner, thereby modulating the process of substrate degradation. The cartoon in Fig. 1A illustrates the structure of 26S proteasome that are bound with USP14. For more details, we recommend referring to references [3,9,27]. As shown in Fig. 1A, USP14 consists of two domains, namely UBL domain and USP domain. The UBL domain resembles the structure of ubiquitin and can bind to the T2 site of RPN1 in the proteasome. The USP domain possesses the capability of deubiquitination and contacts the exterior of the OB-ring opposite RPN11. The OB-ring forms the entrance of the polypeptide substrate into the ATPase motor and CP. The substrate is unfolded and translocated by the ATPase motor, and hydrolyzed within the CP.

**Fig. 1.**
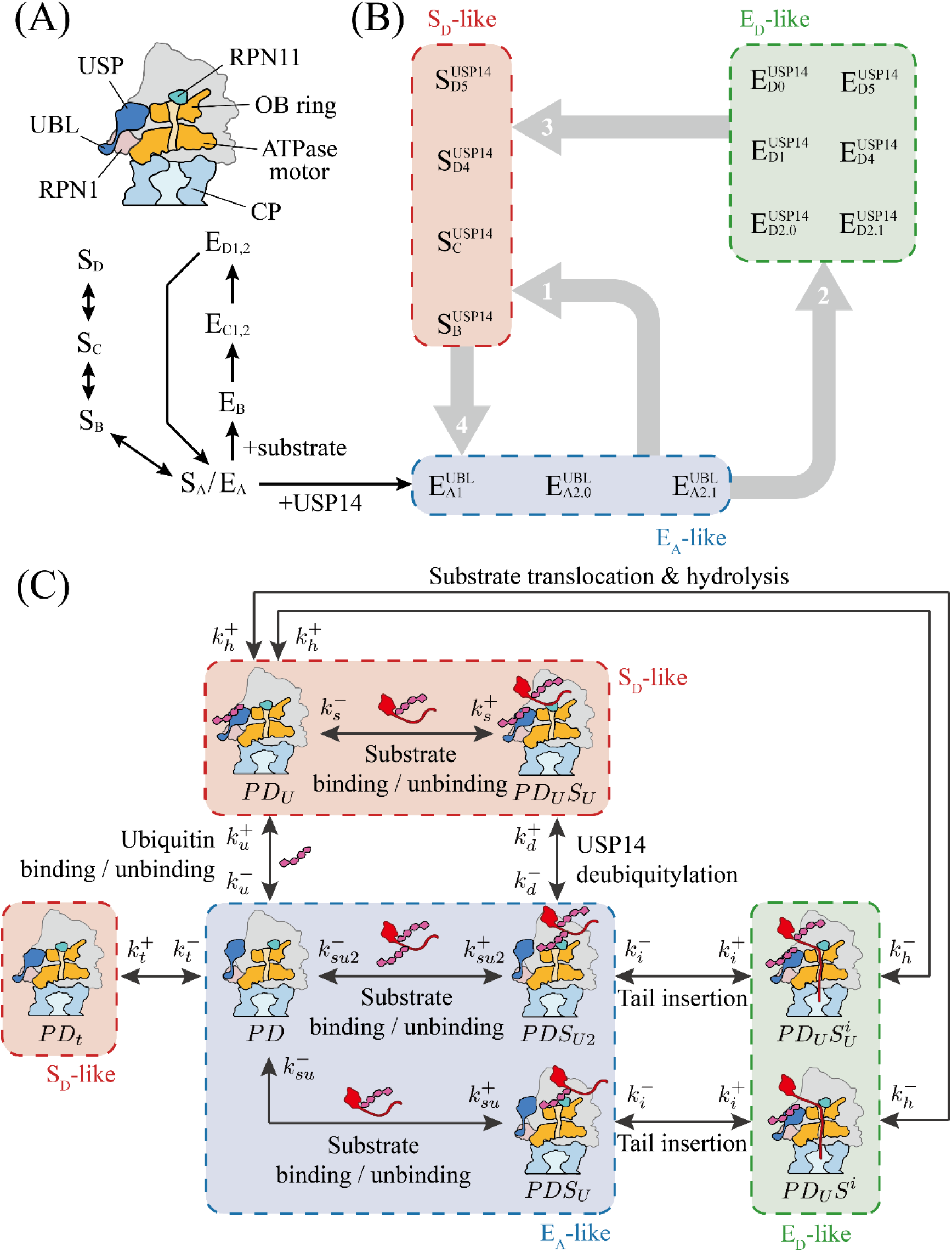
USP14-bound 26S proteasome in substrate degradationn. (A) The structure of USP14-bound proteasome. (B) Conformations of the human 26S proteasome and their transitions known from experiments [3,9,27]. *S*_*A*_. *S*_*B*_. *S*_*C*_. *S*_*D*_ (*E*_*A*_, *E*_*B*_, *E*_*C*1.*C*2_, *E*_*D*1.*D*2_) are for proteasomal conformations free of substrate (in the presence of substrate), with subscripts *A, B, C, D* (and the numbers) representing conformational (and sub-conformational) differences. The E_A_-like, E_D_-like, and S_D_-like conformations shown in dashed boxes are for USP14-bound proteasome, with structures similar to E_A_, E_D_, and S_D_. The superscript *UBL* (or USP14) denotes that the proteasome is bound with the UBL domain (or with both UBL and USP domains of USP14). The thin arrows (or thick gray arrows) are for experimentally certain (or less certain) conformation transitions. (C) The reaction network for USP14-regulated proteasomal substrate degradation reformulated from (B). Constituents of the proteasomal complex are denoted with *P* (for 26S proteasome), *D* (for USP14), and *S* (for substrates), respectively.

Figure 1B summarizes the recent experimental findings on the human 26S proteasomal conformations and transitions during substrate degradation [3,9,27]. Four distinct proteasomal conformations (S_*A*_. *S*_*B*_. *S*_*C*_. *S*_*D*_) in the absence of both substrate and USP14 have been identified [9], and four types of conformations (E_*A*_. E_*B*_. E_*C*1.*C*2_. E_*D*1.*D*2_) in the presence of substrate have been recognized for the USP14-free proteasome [3]. Specifically, the substrate is not yet bound to the proteasome in *E*_*A*_ (identical to S_*A*_), and is bound to the proteasome via the ubiquitin chain in *E*_*B*_ (without its N- or C-terminus being inserted into the ATPase motor of the proteasome). In *E*_*C*_, the substrate is inserted into the ATPase motor and deubiquitinated by RPN11. It is subsequently translocated by the ATPase motor into the CP and hydrolyzed in *E*_*D*_. The S_*A*_. *S*_*B*_. *S*_*C*_. *S*_*D*_ conformations have similar structures to E_*A*_. E_*B*_. E_*C*_. E_*D*_, respectively, despite the absence of substrate [2,9].

For substrate degradation in 26S proteasome that is regulated by USP14, recent time-resolved cryo-EM experiments have disclosed a qualitative picture for proteasomal conformations and their temporal changes [27]. As illustrated with dashed boxes in Fig. 1B, the recently observed 13 conformations of USP14-bound proteasome were categorized into E_A_-like 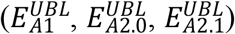 E_D_-like 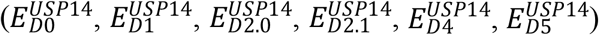, and S_D_-like 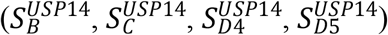 classes. The E_A_-like, E_D_-like, and S_D_-like conformations have structures similar to E_A_, E_D_, and S_D_, which are not regulated by USP14. For more detailed information on the conformational difference in E_A_-like, E_D_-like, and S_D_-like proteasomal states, please refer to the description in Supporting information. As indicated with thick gray links in Fig. 1B, transitions occur among the E_A_-like, S_D_-like, and E_D_-like conformations. Previous studies illustrated that in E_A_-like conformations, competition exists between the deubiquitination of USP14 and the tail insertion of substrate [25]. The substrate could not be stably bound to the proteasome if the substrate ubiquitin is removed by USP14 before the tail insertion completes. This can make E_A_-like states change into the S_D_-like conformations (indicated by gray arrow 1 in Fig. 1B). Conversely, if ubiquitin is cleaved after tail insertion, the substrate can maintain a stable binding due to the force exerted by ATPase motor, facilitating subsequent unfolding and degradation processes, *i*.*e*., E_A_-like changes to E_D_-like (refer to gray arrow 2 in Fig. 1B). After the completion of substrate hydrolysis, the proteasome in E_D_-like conformations returns to the unengaged S_D_-like conformations [29], as denoted by gray arrow 3 in Fig. 1B. Ultimately, the proteasome in the substrate-resistant S_D_-like conformations changes back to the E_A_-like conformations (refer to gray arrow 4 in Fig. 1B).

### 2.2 Reaction network and kinetic model for USP14-regulated proteasomal substrate degradation

Based on the experimental findings as in Fig. 1B, we sought to construct a reaction network for USP14-regulated E_A_-like, E_D_-like, or S_D_-like conformations in substrate degradation (Fig. 1C). In our model, the 13 distinct conformations are not considered directly as variables but are classified according to their differences and changes in the constituents of the complex. To account for conformational changes of the proteasome, the primary components of the complex are represented by the symbols *P* for the 26S proteasome, *D* for USP14, and *S* for substrates. For instances, the combination *PD* represents the 26S proteasome bounded with USP14, and *PDS* denotes the proteasome-USP14-substrate complex. For simplicity, the conformational differences in the ATPase motor are ignored in our model, and the proteasomal complex is characterized by whether it is bound with the ubiquitin chain (denoted by subscript *U*) and whether the substrate is engaged with its initial tail inserted into the OB-ring of the proteasome (denoted by superscript *i*). For an instance, *PD*_*U*_ represents the proteasome-USP14 complex with USP14 bound with the ubiquitin chain. Partially deubiquitination is permitted in our model. *S*_*U*2_ represents the substrate not deubiquitinated by USP14, and partially deubiquitinated substrate by USP14 is denoted with *S*_*U*_. Applying the above scheme, the E_A_-like conformations 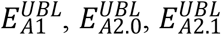 involved in the processes before tail insertion or deubiquitination are represented with *PD, PDS*_*U*2_, *PDS*_*U*_ in our model. The six E_D_-like conformations responsible for substrate translocation and degradation are casted briefly in two states 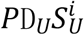 and *PD*_*U*_*S*^*i*^. Similarly, the four S_D_-like conformations, involved in translocation inhibition, are simplified into two states *PD*_*U*_ *S*_*U*_ and *PD*_*U*_. In Fig. 1C, *PD*_*t*_ represents a state before the addition of substrate. It belongs to the S_D_-like class which undergoes a reversible transition to the *PD* state in E_A_-like type conformations in absence of substrate.

To take into account the experimental findings of conformational changes as illustrated in Fig. 1B, state transitions occur in the simplified states in our model (refer to Fig. 1C). Substrate binding/unbinding take place in the E_A_-like states (*PD, PDS*_*U*2_, *PDS*_*U*_) and in the S_D_-like states (*PD*_*U*_*S*_*U*_, *PD*_*U*_), thus cause transitions between the E_A_-like and S_D_-like states. Secondly, the E_A_-like states *PDS*_*U*2_, *PDS*_*U*_ can change into E_D_-like states 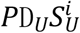 and *PD*_*U*_*S*^*i*^ by tail insertion of substrate engagement. The accomplishment of substrate translocation and hydrolysis transforms the E_D_-like states 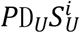 and *PD*_*U*_*S*^*i*^ into the S_D_-like state *PD*_*U*_. In addition, the initial E_A_-like state *PD* (or *PDS*_*U*2_) can be converted to the S_D_-like state *PD*_*U*_ (or *PD*_*U*_*S*_*U*_) by ubiquitin binding (or deubiquitylation of USP14).

In our model, we assume that 26S proteasomes, USP14, and substrates are present in abundance and the reactions among them follow the mass action law. The chemical reaction dynamics for substrate degradation in 26S proteasome regulated by USP14 in Fig. 1C can be described with eleven variables, eight of which represent the concentrations of E_A_-like states (*PD, PDS*_*U*2_, *PDS*_*U*_), E_D_-like 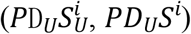, and S_D_-like (*PD*_*U*_*S*_*U*_, *PD*_*U*_, *PD*_*t*_) states for USP14-bound proteasome, two for the concentrations of substrates (*S*_*U*2_, *S*_*U*_), and one for the concentration of ubiquitin chain (*U*). The ordinary differential equations for the reactions are listed in Supporting information (Eqs. S1).

### 2.3 The dynamics of USP14-bound proteasomal substrate degradation

The coupled ordinary differential equations (Eqs. S1) for the kinetic model of USP14-regulated proteasomal substrate degradation have been numerically solved, and the results are presented in Fig. 2. The time evolution in the proportions of E_A_-like (*PD, PDS*_*U*2_, *PDS*_*U*_), E_D_-like 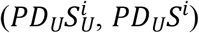, and S_D_-like (*PD S, PD, PD*) conformations is readily obtained from the simulated dynamics of eight variables for USP14-bound proteasome concentrations, as illustrated in Fig. 3A. The temporal change in the ratio of the residual substrate concentration (*S*_*U*2_, *S*_*U*_) relative to the initial amount is also shown. Upon substrate addition at zero minute, the proportion of E_D_-like conformations, which are involved in substrate engagement, translocation, and hydrolysis, increases rapidly in the first minute, and falls slowly from the peak value in the following thirty minutes. On the contrary, the proportion of E_A_-like conformations, which are responsible for the processes before substrate tail insertion or deubiquitination, decreases rapidly to the bottom value and recovers slowly as the substrate is degraded. The two opposing trends arises naturally due to the conservation in the total proteasome concentration. The proportion of S_D_-like conformations, which are associated with degradation inhibition mediated by ubiquitin-bound USP14, maintains at low percentages and has no prominent changes during degradation as the residual substrate concentration decreases continuously. At 30 minutes, there are approximately 25% of the substrate remained undegraded. Recent time-resolved cryo-electron microscopy (cryo-EM) experiments have elucidated the dynamic shifts in the proportions of 13 distinct conformational states as well as the progressive decrease in residual substrate levels over time [27]. These experimental findings were obtained at a temperature of 10°C under specific conditions: 1 mM ATP, 10 μM substrate Sic1^PY^, and 1 μM USP14-bound proteasome. As illustrated in Fig. 3A, the model results (curves) agree closely with the experimental findings (the dots).

**Fig. 2.**
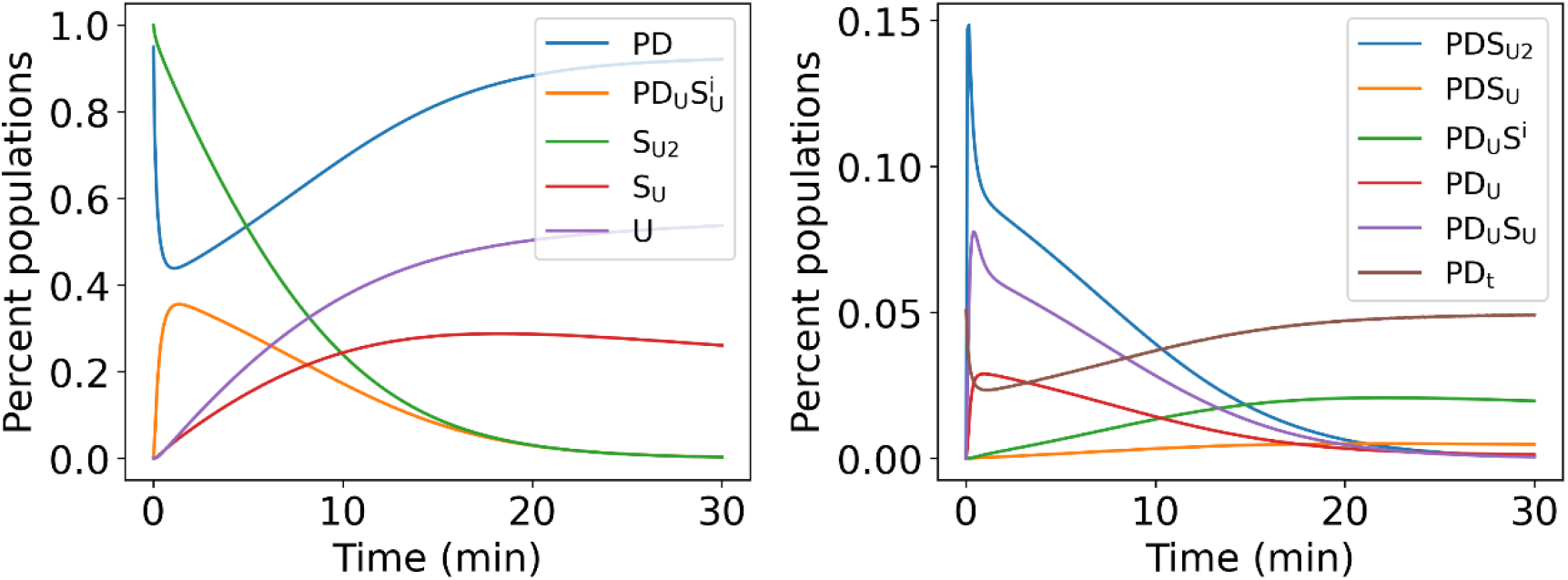
Time evolution for the reactions in Fig. 1C for USP14-regulated proteasomal substrate degradation. The results are obtained by numerical simulations of Eqs. S1. Detailed information of calculation is given in Method. The parameters used in simulation are listed in Supporting information (Table S1).

**Fig. 3.**
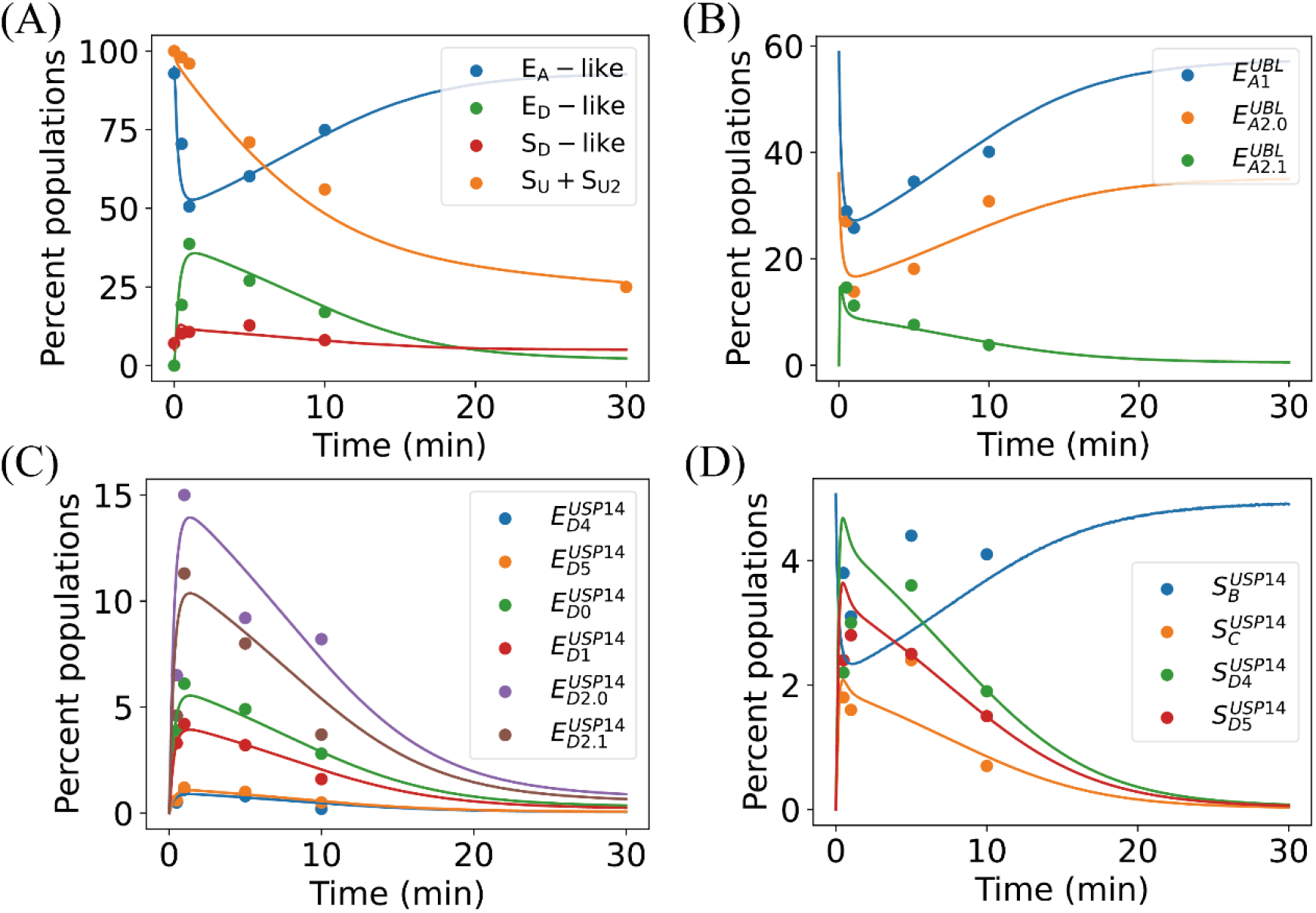
Dynamics of the USP14-bound proteasome. (A) Time evolution of the proportions of E_A_-like, E_D_-like, and S_D_-like categories of conformation, as well as the concentration of residual substrate (*S*_*U*2_. *S*_*U*_) relative to the initial amount over time. (B) Temporal changes in the proportions of 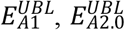, and 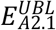 conformations within the E_A_-like conformation. (C) Temporal changes in the proportions of 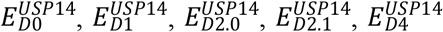, and 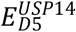 conformations within the E_D_-like conformation. (D) Temporal changes in the proportions of 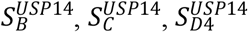, and 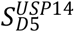 conformations within the S_D_-like conformation. Solid lines represent simulation results, and dots are from the experimental data [27].

As the 13 intermediate conformations are not explicitly included in our model as dynamical variables (due to lack of detailed experimental information of conformational changes), the proportions and their dynamical changes cannot be directly obtained in our simulations. Based on reasonable assumptions, the experimentally observed changes in the proportions of 13 intermediate conformations can still be inferred in our model. Firstly, the six E_D_-like conformations (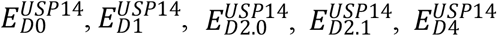, and 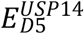) are involved primarily in fast changes of the ATPase motor during substrate unfolding and translocations. As they represent a series of intermediates changing much faster than the complete rounds of substrate translocation, the transitions between the six E_D_-like conformations are effectively in quasi-equilibrium. It is reasonable to assume that the proportions of these six intermediates in the whole of E_D_-like conformations are fixed during the substrate degradation. In our simulations, the fixed percentages are obtained by fitting the experimental results, which are 2.5% for 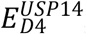, 3.0% for 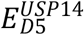, 15.5% for 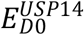, 11.0% for 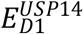, 39.0% for 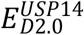, and 29.0% for 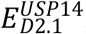, respectively. As shown in Fig. 3C, the dynamical changes in the proportions of the six E_D_-like conformations obtained from our simulations closely match the experimental data.

For the three E_A_-like conformations, 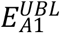 lacks any visible substrate, 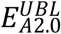 shows ubiquitin bound at the RPN11, and 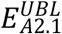 exhibits both ubiquitin and subtle substrate density on RPN11. Given the difficulty of cryo-EM in visualizing substrate conformations outside the OB-ring, we consider 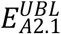 to encompass both *PDS*_*U*2_ and *PDS*_*U*_ states in our model, *i*.*e*., 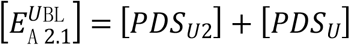. We hypothesize that 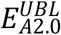 represents an unstable binding of ubiquitin (possibly originated in substrate-tagged ubiquitin chains or ubiquitin chains generated by USP14 deubiquitination). Considering the constant total number of ubiquitin chains in experiment, we assume that 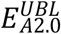 maintains a rapid equilibrium with 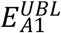 and corresponds concurrently to the *PD* state in the model. Thus, the percentages of 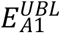 and 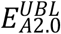 in the whole of *PD* conformation can be considered as fixed, which are similarly fitted to be 62.0% and 38.0%, respectively. The results in Fig. 3B show that the model results are in good agreement with the experimental data.

The four S_D_-like conformations primarily differ in the status of the *CP* gate and the conformation of the ATPase motor, making it challenging to establish a direct correspondence to our model based solely on their conformational features. In our simulations, we hypothesize that 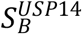 directly corresponds to the *PD*_*t*_ state (*i*.*e*.,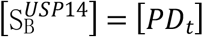), and that 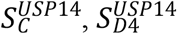, and 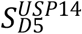 correspond to the *PD* _*U*_*S*_*U*_ and *PD*_*U*_ states. Due to the reduced affinity between the substrate and the proteasome after USP14-mediated deubiquitination, cryo-EM images are likely unable to differentiate the *PD*_*U*_*S*_*U*_ and *PD*_*U*_ states. The transitions among 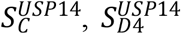, and 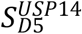 may achieve rapid equilibrium. Their percentages in the whole of *PD*_*U*_*S*_*U*_ and *PD*_*U*_ are fixed, which have been fitted to be 20.5% 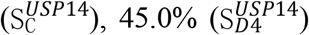, and 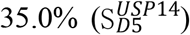, respectively. In Fig. 3D, the simulation results are basically consistent with the experimental data. The discrepancy might be due to the relatively low proportions of S_D_-like conformations in the total of 13 conformations and that experimental measurements of their proportions may be more susceptible to errors in measurements. The simulation results in Fig. 3 suggest that the kinetic model adequately explain the experimental observations known up to date.

## 3. Simplified model and the influence of USP14, substrate and ATP on substrate degradation

### 3.1 The effect of USP14 on substrate degradation rate

To check into the influence of USP14 on substrate degradation by the proteasome, we proceed with additional simplifications to the kinetic model that we have discussed above. Specifically, we neglect the more detailed *PDS*_*U*_ and 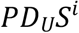 states in Fig. 1C and consider only the case where substrate not deubiquitinated by USP14 binds to the proteasome. In considering the homeostasis within the cell, we assume that the concentrations of *S*_*U*2_, *S*_*U*_, and *U* remain constant. In Fig. 4A, the reaction network for USP14-regulated proteasomal substrate degradation in Fig. 1C is simplified as the substrate degradation loop 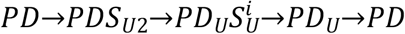 coupled with the loop of deubiquitination *PD*→*PDS*_*U*2_→*PD*_*U*_*S*_*U*_→*PD*_*U*_→*PD*. In comparison, Fig. 4B illustrates a simple loop of substrate degradation 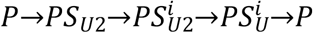, in which the proteasome is not regulated by USP14. The kinetic equations of mass action law are assumed for the networks in Fig. 4A and Fig. 4B and are given in Supporting information (Eqs. S2 and S3) together with the parameters (Tabs. S2 and S3).

**Fig. 4.**
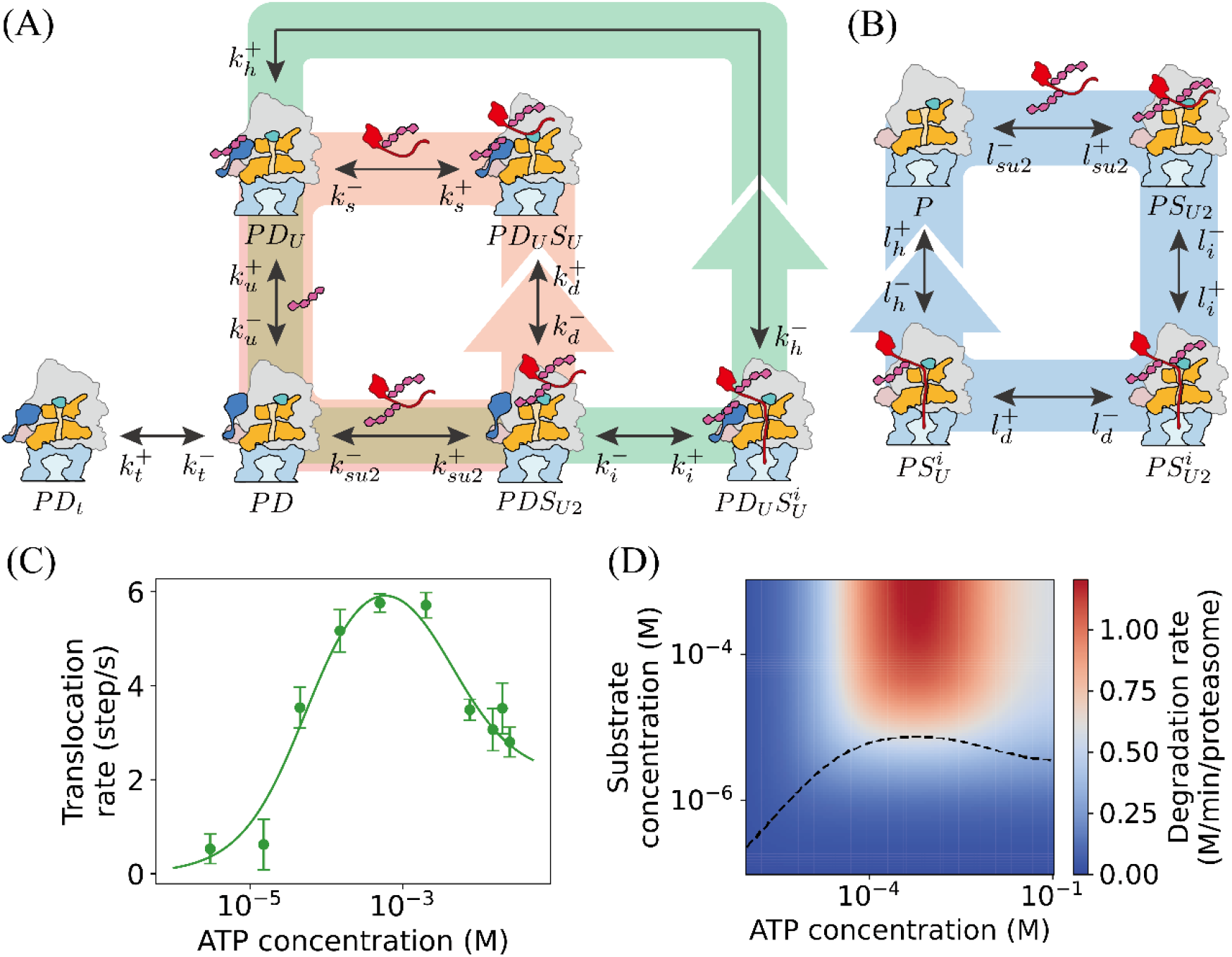
Simplified model and the influence of substrate and ATP on substrate degradation. (A) Simplification of the reaction network in Fig. 1C for USP14-regulated proteasomal substrate degradation. Pink and green arrows represent the reaction loop for USP14-mediated deubiquitination and the loop for substrate degradation, respectively. (B) A simple reaction network of proteasomal substrate degradation free of USP14. Blue arrows indicate the loop of substrate degradation. (C) The experimentally observed substrate translocation rates (dots) fitted with the formular of Eq. 5 (curve). (D) The dependence of substrate degradation rate on the concentrations of substrate and ATP predicted by Eqs. 3, 5, and 6. The black dashed line denotes the half-maximal effective concentration (EC_50_) of the substrate under various ATP concentrations. The parameter values used are listed in Tab. S2.

The influence of USP14 on the rate of proteasomal substrate degradation can be obtained by analyzing the simplified model for USP14-regulated substrate degradation (Fig. 4A) in comparison with the simple reaction model for the degradation without USP14 (Fig. 4B). For both cases, the steady-state substrate degradation rates can be obtained analytically by setting the right-hand sides of Eqs. S2 and Eqs. S3 to zero. In Fig. 4B, the processes of substrate insertion, translocation and hydrolysis, deubiquitination, unbinding of ubiquitin and deubiquitinated substrate are nearly irreversible, the relevant parameters in Tab. S3 are negligible with 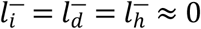. The substrate degradation rate *v*_―*USP*14_ for the USP14-free process can be deduced to be,

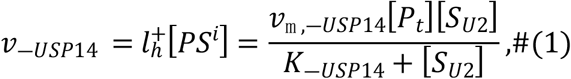

where 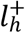 is the rate of substrate hydrolysis, [*P*_*t*_] is the total concentration of all proteasomal states, 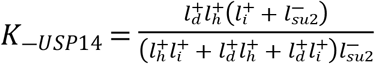, and *v*_*m*.―*USP*14_ is the saturated rate,

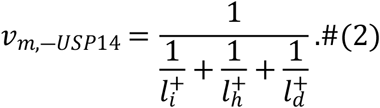

Similarly, for the case of substrate degradation regulated by USP14 (Fig. 4A), the relevant rate parameters in Tab. S2 for the negligible inverse reactions are assumed as zero with 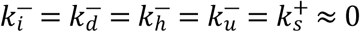, the substrate degradation rate *v*_+*USP*14_, which is 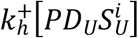, for USP14-regulated proteasomal degradation rate takes the form,

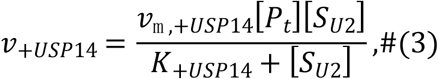

in which, the saturated degradation rate is,

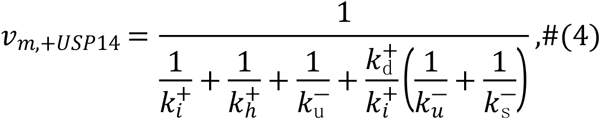

and 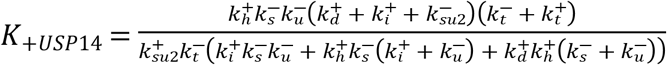

The effect of USP14 on degradation can be obtained by checking the substrate degradation rates of Eq. 3 in comparison with Eq. 1. As the expressions of Eqs. 1 and 3 are complex, it is not convenient to evaluate the relative size of *v*_―*USP*14_ and *v*_+*USP*14_ to determine whether the influence of USP14 on the substrate degradation rate is positive or not. Take a step back, the saturated degradation rates *v*_*m*.―*USP*14_ and *v*_m .+*USP*14_ of Eqs. 2 and 4 can be compared due to their simpler forms. Typically, the processes of substrate insertion and hydrolysis are not influenced by the binding of USP14, thus 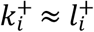 and 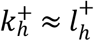. The parameter 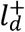 in Eq. 2 is the deubiquitination rate constant of RPN11 in the absence of USP14, and 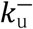 in Eq. 4 is the rate constant for ubiquitin falling off USP14 in the presence of USP14. According to recent experiments, 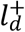 is from one to two orders of magnitude larger than 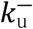 [7]. As 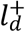 in Eq. 2 is generally much greater than 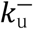 in Eq. 4, it is always the case that *v*_.+*USP*14_ < *v*_*m*.―*USP*14_. Therefore, under conditions of substrate saturation, the typical effect of USP14 regulation is to reduce the rate at which substrates are degraded by the proteasome. The model result is consistent with the experimental findings that demonstrated USP14 inhibits the substrate degradation by the proteasome [23,25,27].

### 3.2 The dependency of the degradation rate on substrate and ATP concentrations for USP14-bound proteasome

The rate *v*_+*USP*14_ in Eq. 3 for USP14-regulated proteasomal protein degradation not only depends on the concentration of substrate [*S*_*U*2_], but also a function of the substrate hydrolysis rate 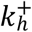 which is determined by the concentration of ATP. Previous biochemical experiments have highlighted that the rate of substrate translocation in the proteasome doesn’t rise monotonically with increasing ATP concentration [30]. As depicted in Fig. 4C (dots), the rate follows a biphasic pattern. Initially, the rate increases as ATP concentration goes up, then paradoxically decreases at higher ATP levels. The experimental substrate translocation rate in (30) is found to be best fitted with the following Hill functional form (curve in Fig. 4C):

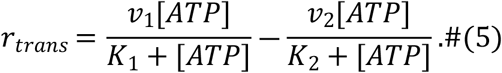

Where *v*_1_ = 7.214 step s, *v*_2_ = 5.165 step s, *K*_1_ = 5.666 × 10^―5^ M, and *K*_2_ = 3.962 × 10^―3^ M. The rate of substrate hydrolysis depends linearly on the translocation rate,

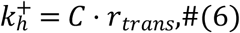

where *C* is the coefficient. It is estimated here to be 0.307(M · s) (min · step) from the experimentally observed translocation rate in (30) under conditions of 1mM ATP and the hydrolysis rate 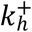 we use in our simulations of Eqs. S1.

The dependence of the USP14-regulated proteasomal hydrolysis rate on both substrate and ATP concentrations can be readily obtained by substituting Eq. 6 into Eq. 3. Figure 4D depicts the substrate degradation rate *v*_+*USP*14_ as the function of [*S*_*U*2_] and [*ATP*] concentrations with the parameter values in Tab. S2. The peak of the translocation rate does not appear in the upper-right corner of [*S*_*U*2_]-[*ATP*] space due to the biphasic relationship between the substrate translocation rate and ATP concentration. From Fig. 4D, the range of ATP concentration for optimal degradation remains consistent for different substrate concentrations larger than 0.01 mM. Also illustrated in Fig. 4D is the black dashed line denoting the half-maximal effective concentration (EC50) of the substrate across different ATP concentrations. EC50 is defined as the substrate concentration corresponding to half the maximum degradation rate at each ATP concentration. It rises as the ATP concentration increases, and then declines once the optimal ATP concentration has been exceeded. The EC50 curve represents the interplay between the ATPase motor’s dynamics and other processes occurring in the proteasome bound by USP14, a relationship that can be verified through future experimental studies.

### 3.3 The influence of USP14 on substrate preference in proteasomal degradation

Within living cells, proteasomes frequently encounter the task of concurrently degrading multiple types of substrates. Given the distinct properties of these substrates, the degradation rates often differ significantly, and proteasomes exhibit preferences in the degradation of substrates [31]. The interplay between proteasomes and USP14 modulates the kinetics of substrate degradation, thereby would shape the selective preferences of the 26S proteasome towards various substrates. To investigate the influence of USP14 on substrate selectivity, we consider that two different substrates, denoted as *S* and *T*, are present in the reactions of Fig. 4A and Fig. 4B. The reaction kinetic equations, for both with and without the regulation of USP14, are detailed in Supporting information (Eqs. S4 and Eqs. S5). The degradation preference of substrate *S* over substrate *T* can be quantified by the ratio of their respective degradation rates., *i*.*e*., *P*_*ST*_ = *v*_*S*_/*v*_*T*_. The influence of USP14 on the substrate preference during degradation can be estimated by comparing the relative magnitudes of the preferences *P*_*ST*.+*USP*14_ in the presence of USP14 and *P*_*ST*.―*USP*14_ in its absence. It is reasonable to assume that USP14 would not influence substrate binding/unbinding and tail insertion into the proteasome, and the relevant rate constants for substrate *S* (*k*’s) and substrate *T* (*l*’s) are approximately equal (refer to Method). On this basis, the preference ratio *η* ≡ *P*_*ST*.+*USP*14_/*P*_ST.―*USP*14_ at the stationary state is found to be,

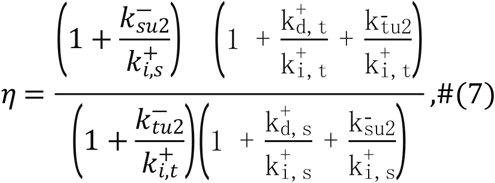

where 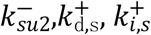 (and 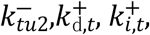.) are the rate constant of substrate unbinding, deubiquitination of USP14, and substrate insertion into the ATPase motor for *S* (and *T*). The rate constants of unbinding 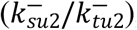, deubiquitination 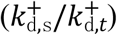, and insertion 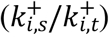 for substrate *S* relative to substrate *T* determine the preference ratio *η* in combination. For USP14 to enhance the degradation preference of substrate *S*, it is necessary that *η* > 1 or, equivalently.

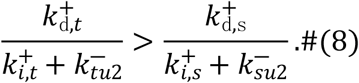

In case *η* = 1, the above inequality holds with equality. The influence of USP14, quantified by Eq. 7, on the preferential degradation of substrate *S* over *T* is vividly depicted through the heat maps in Fig 5. The maps illustrate the relationship in the relative rate space 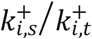 versus 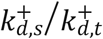 (Fig. 5A), and further in the alternative relative rate space 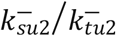 plotted against 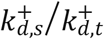 (Fig. 5B). The intermediate dividing line with *η* = 1 is marked. From Eqs. 7 and 8 and Fig. 5, it can be concluded that with relatively large ratios of 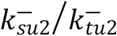 or 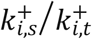, or small 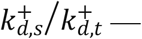 in other words, when the substrate is easier to dissociate from the proteasome, or readily inserted into the OB-ring, or less sensitive to ubiquitination — its proteasomal degradation would be enhanced by the binding of USP14 to the proteasome.

**Fig. 5.**
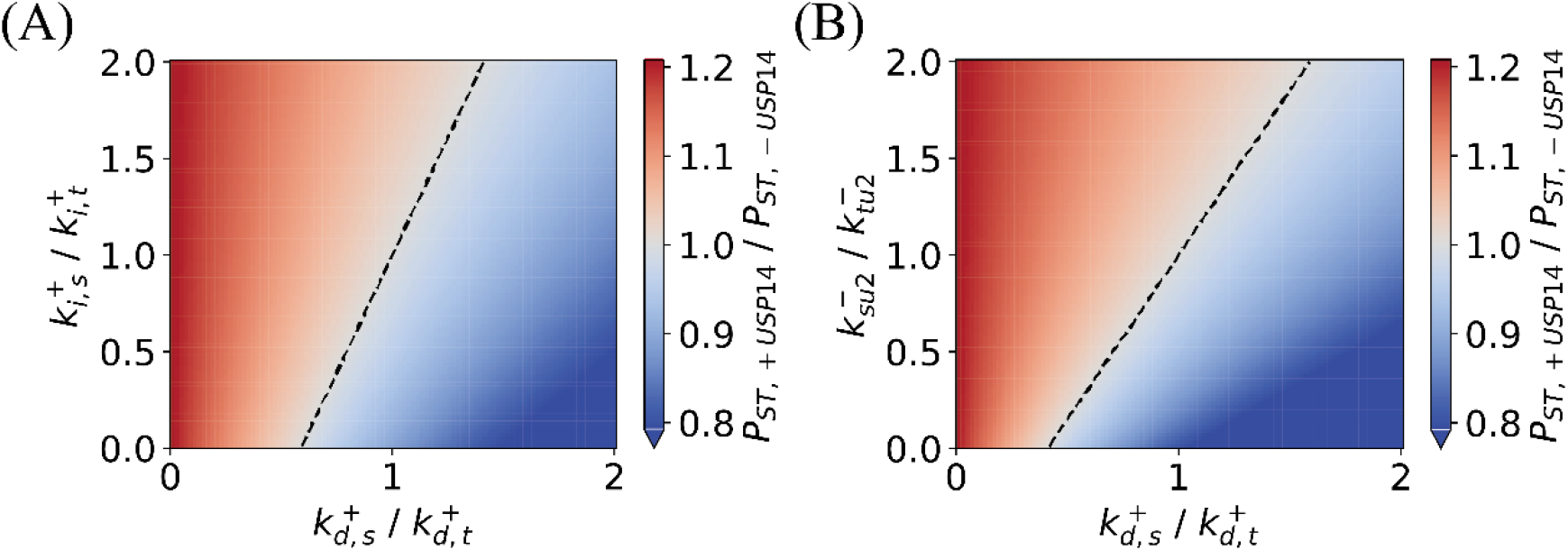
Effects of USP14 on substrate preference in proteasomal degradation. The preference ratio η in Eq. 8 is illustrated in the relative rate space 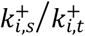 versus 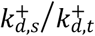 (A), and in 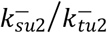 versus 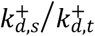 space (B). The black dashed line indicates *η* = 1 where the degradation preferences of substrate *S* to *T* in USP14-bound and USP14-free proteasomes are equal. Parameters: 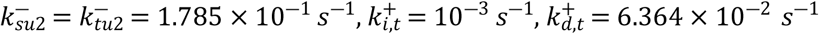 for (A), and 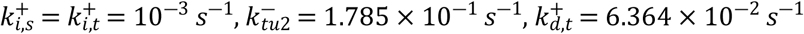 for (B).

## 4. Discussion

The ubiquitin-proteasome system (UPS), which includes USP14 and the 26S proteasome, is crucial for the regulation of protein turnover in cells. Studying their interaction provides insight into the mechanisms underlying protein degradation, which is fundamental to cellular homeostasis. The process of proteasome-mediated protein degradation occurs rapidly, with timescales on the order of milliseconds to seconds. Capturing intermediate states during this process is technically challenging due to the fast kinetics involved. The addition of USP14 introduces further complexity to its conformational changes. Distinguishing and classifying these dynamic states at high resolution requires sophisticated cryo-EM imaging analysis and classification algorithms. Up to now, time-resolved cryo-EM experiments on USP14-regulated allostery of the 26S proteasome are very limited, and an accurate understanding of the highly dynamic machine in substrate degradation remains elusive [27]. For instances, cryo-EM experiments have provided only fragmentary and vague views on the rapid conformational changes of the ATPase motor, and it is difficult to obtain cryo-EM images for substrate conformations outside the OB-ring due to its structural flexibility under thermal noise.

The limitations in experimental studies necessitate the use of mathematical modeling to hypothesize in experimental uncertainties and to gain complementary insights into the intricate substrate degradation process. This is particularly pertinent when detailed cryo-EM studies, aiming to capture time-resolved proteasomal allostery, are challenging to achieve. In this paper, we presented a kinetic model which is based on recent experimental findings in USP14-bound proteasomal conformations in substrate degradation. Our model accurately interpreted the recent experimental data on the time-resolved variations in the conformational landscape of human 26S proteasomes, as detailed in the study of [27]. Consistent with experimental findings, the model demonstrated that the proteasome’s rate of substrate degradation is reduced under the regulatory influence of USP14. Our model illustrated how the rate of degradation is influenced by the concentrations of substrate and ATP, and it predicted the EC50 value of the substrate for varying ATP concentrations. When confronted with multiple substrates, USP14 influences the proteasome’s preference, leading it to favor substrates that can readily engage with the OB-ring, are weaker bound to the proteasome, and are less susceptible to deubiquitination processes.

The regulatory role of USP14 on the proteasome has been a focus in study of the USP14-proteasome complex. Several studies have demonstrated its inhibitory effect on substrate degradation [23,25,27]. It is important to note that the present experimental evidence is not comprehensive enough to definitively ascertain the inhibitory role of USP14 across the entire spectrum of substrates. Our model suggested that the inhibitory effect of USP14 depends on several parameters that are rooted in the structural characteristics of the substrate. Theoretically, there might exist a class of substrates upon which USP14 could have a facilitative impact. In fact, USP14 has been recently found to exert activating effects on the proteasome [32]. It was reported that if the proteasome binds to USP14 with only the UBL domain, it can up-regulate the degradation rate of substrates by the proteasome. The precise mechanism underlying the heightened degradation rate observed with the UBL-domain-only USP14 remains unclear. As such an incomplete USP14 does not exist under *in vivo* conditions, the observation may not apply to the case of complete USP14 with both UBL and USP domains. In fact, the absence of the USP domain in the truncated form prevents USP14 from executing its deubiquitinating function on substrates. The regulation of USP14 without the USP domain to the 26S proteasome might be very different from the situation we considered here. To date, there have been no experimental reports indicating that complete USP14 can achieve an increase in degradation rates.

Experimental findings regarding substrate preference and selectivity of the 26S proteasome in substrate degradation have been accumulated. Several factors have been revealed that can influence the substrate selectivity, including the amino acid sequence and the length of the initiation region of substrate [33–36], the number of ubiquitin chains, as well as the position, length, and type of each ubiquitin chain [31,37]. The specific role of USP14 in altering proteasomal substrate selectivity is still unclear. Our model analyses suggest that a substrate, when compared to others, which is more readily inserted into the OB-ring and easily disengaged from the proteasome, while simultaneously being less susceptible to deubiquitination by USP14, would be preferentially targeted for degradation by the proteasome complexed with USP14. This provides a fresh perspective on the potential biological roles of USP14 and encourages further experimental exploration to better understand its functions.

## 5. Method

In the simulations of Eqs. S1, the ordinary differential equations are numerically solved and analyzed using Python SciPy (version 1.7.3) and NumPy library (version 1.22.4). We assume that before the addition of substrate, the transition between the USP14-bound proteasome *PD* state and the *PD*_*t*_ state, in which the entry to the OB-ring is blocked by RPN11, is at equilibrium. In accordance with the experimental setup [27], we set the initial concentrations for *PD, PD*_*t*_, and *S*_*U*2_ as 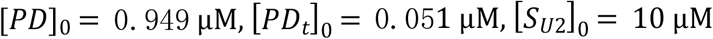, while all other variables are initialized to zero. The rate constants for substrate binding and dissociation from the proteasome, as well as those for substrate insertion and translocation, along with the rate of substrate hydrolysis in the ODEs of Eqs. S1, are adopted from experimental measurements (6,30,31). Since the experiments were conducted at temperatures higher than those of the time-resolved cryo-EM experiments (below 10 °C), we use the experimentally measured rate parameters as upper limits in our model. The remaining rate constants were determined by fitting the simulation results to the time-resolved cryo-EM experimental data, as shown in Fig. 3A. The full parameter values for Eqs. 1 are listed in Supporting information (Tab. S1). We also conduct a parameter sensitivity analysis (Fig. S1 in Supporting information), which highlights the critical roles of the substrate binding rate, tail insertion rate, deubiquitination rate, and substrate translocation hydrolysis rate in USP14-mediated regulation of proteasomal processes. In our simulations of the temporal changes in the percentage populations of 13 intermediate conformations of the USP14-bound proteasome (shown in Fig. 3B, 3C, and 3D), we assume that quasi-equilibriums have been achieved within the E_A_-like, E_D_-like, and S_D_-like classes during substrate degradation. While the concentrations in each class undergo temporal changes, the ratios between them should remain roughly constant. In our simulations, the fixed ratios are determined by fitting the model results to the experimental observations.

To evaluate the effects of USP14 on proteasomal degradation and substrate preferences, we adopt the simplified model in Fig. 4A and compare it to the scenario depicted in Fig. 4B, which lacks USP14. The kinetic equations for the reaction networks in Fig. 4A and Fig. 4B are listed in Supporting information (Eqs. S2 and Eqs. S3), with rate constants denoted differently by *k*’s and *l*’s, respectively. For both USP14-free and USP14-bound proteasomes, we use identical concentrations of proteasome, substrate, and ATP at the same temperature in our analyses to compare the degradation processes.

We also assume that the influences of USP14 on substrate binding and release, tail insertion, and translocation and hydrolysis rates can be disregard, *i*.*e*., 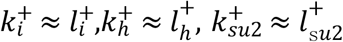 in considering the effect of USP14 on proteasome degradation rates. Similarly, when assessing the effect of USP14 on proteasome substrate preference, 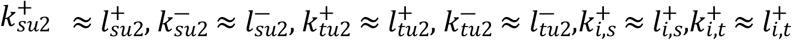 are assumed.

The expression of *η* (Eq. 7) is derived from the steady-state of Eqs. 4 and Eqs. 5. The preference of substrate *S* over substrate *T* in USP14-regulated proteasomal degradation, which is calculated as 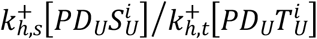,is determined and given by,

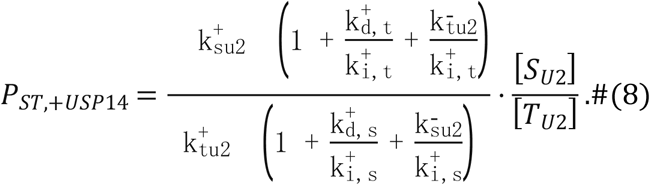

For the case of USP14-free degradation, the preference, which is equal to 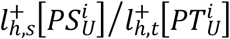, has the form,

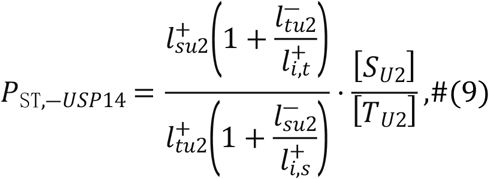

where 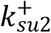 and 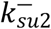 (and 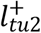 and 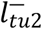) are the rate constants for binding and unbinding of ubiquitin-tagged substrate *S*_*U*2_ (and *T*_*U*2_) to the proteasome, with subscripts ‘s’ and ‘t’ for substrate *S* and substrate *T*, respectively. The magnitudes of *P*_*ST*.+*USP*14_ and *P*_*ST*.―*USP*14_ in Eqs. 8 and 9 depend on the parameters *k*’s and *l*’s. The ratio of Eq. 8 and Eq. 9 gives the equation for *η* (Eq. 7). Inequality Eq. 8 is deduced from expression Eq. 7 by letting *η* > 1.

## Supporting information

**Text S1**. Conformational differences in E_A_-like, E_D_-like, and S_D_-like proteasomal states.

**Table S1**. The parameters in Eqs. S1 for the full reaction model in Fig. 1C.

**Table S2**. The parameters in Eqs. S2 for the simplified model for USP14-regulated proteasomal degradation demonstrated in Fig. 4A.

**Table S3**. The parameters in Eqs. S3 for the reactions of substrate degradation without USP14 in Fig. 4B.

**Equations S1**. ODEs for the simplified kinetic model in Fig. 1C.

**Equations S2**. ODEs for the simplified kinetic model in Fig. 4A.

**Equations S3**. ODEs for the reactions of substrate degradation without USP14 as shown in Fig. 4B.

**Equations S4**. ODEs for USP14-bound and USP14-free proteasome with two substrates *S* and *T*.

**Equations S5**. ODEs for USP14-free proteasome and in the presence of substrates *S* and *T*,

**Figure S1.** Parameter sensitivity analysis for the dynamics of USP14-bound proteasome described by Eqs. S1.

## Acknowledgments

This work was supported in part by the National Natural Science Foundation of China (Grant No. 12090051 and No. 12125401), the National Key Research and Development Program of China (2023YFF1204400 and 2023YFF1204401), the Starry Night Science Fund of Zhejiang University Shanghai Institute for Advanced Study, and AI for Science (AI4S)-Preferred Program, Peking University Shenzhen Graduate School.

## Author Contributions

**Conceptualization**: Youdong Mao, Hongli Wang, Qi Ouyang.

**Formal analysis**: Di Wu, Hongli Wang.

**Funding acquisition**: Hongli Wang, Youdong Mao, Qi Ouyang.

**Investigation**: Di Wu.

**Methodology**: Di Wu, Hongli Wang, Youdong Mao, Qi Ouyang.

**Software**: Di Wu.

**Supervision**: Hongli Wang, Youdong Mao, Qi Ouyang

**Validation**: Di Wu, Hongli Wang.

**Visualization**: Di Wu.

**Writing – original draft**: Di Wu.

**Writing – review & editing**: Hongli Wang, Di Wu.

## Supporting information

### Text S1. Conformational differences in E_A_-like, E_D_-like, and S_D_-like proteasomal states

In E_A_-like conformations, the UBL domain of USP14 tightly binds to the proteasome, while its USP domain has only a weaker interaction with the proteasome. Additionally, substrates are not yet engaged in these conformations, either without substrate 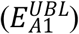 or with substrate loosely bound to the proteasome via ubiquitin chains without insertion of its N- or C-terminus into the ATPase motor 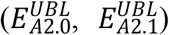. In the E_D_-like conformations, both the UBL and USP domains in USP14 bind to the proteasome, and the substrate ubiquitin chains are also bound to the USP domain. The E_D_-like conformations are engaged with the substrate being gripped into the OB-ring and ATPase motor, and implicated in substrate unfolding, translocation, and hydrolysis. The conformations inside the E_D_-like category differ primarily in their spatial arrangements of the regulatory particle ATPase (RPT) subunits comprising the ATPase motor, as indicated by different subscripts in these conformations. The S_D_-like conformations are not engaged with substrate and are associated with degradation inhibition mediated by ubiquitin-bound USP14. In this case, the USP domain of USP14 remains tightly bound to the proteasome, with ubiquitin chains contact on the USP domain. Moreover, RPN11 blocks the entry to the OB-ring, hindering subsequent substrate insertion and translocation processes. The S_D_-like conformations slightly differ from each other in the states of ATPase motor and the CP gate (the part of CP controlling substrate entrance), and the state of ATPase-CP interface. It is highly likely that transitions occur among the conformations within E_A_-like, E_D_-like, and S_D_-like categories. The exact transitions within the three categories of conformations are still unclear due to the complexity and subtlety of proteasomal machinery.

**Table S1.** The parameters in Eqs. S1 for the full reaction model in Fig. 1C.

Fig. 1C
| parameter | corresponding process | value |
| --- | --- | --- |
| $k_{su2}^+$ | $S_{U2}$ binding on $PD$ | $8.500 \times 10^3 M^{-1}s^{-1}$ |
| $k_{su2}^-$ | $S_{U2}$ unbinding from $PDS_{U2}$ | $1.785 \times 10^{-1} s^{-1}$ |
| $k_{su}^+$ | $S_U$ binding on $PD$ | $5.950 \times 10^2 M^{-1}s^{-1}$ |
| $k_{su}^-$ | $S_U$ unbinding from $PDS_U$ | $1.785 \times 10^{-1} s^{-1}$ |
| $k_i^+$ | tail insertion | $1.250 \times 10^{-1} s^{-1}$ |
| $k_i^-$ | reverse process of tail insertion | $1.000 \times 10^{-3} s^{-1}$ |
| $k_d^+$ | deubiquitination by USP14 | $6.364 \times 10^{-2} s^{-1}$ |
| $k_d^-$ | reverse process of deubiquitination by USP14 | $1.000 \times 10^{-3} s^{-1}$ |
| $k_h^+$ | substrate translocation and hydrolysis | $2.968 \times 10^{-2} s^{-1}$ |
| $k_h^-$ | reverse process of substrate translocation and hydrolysis | $1.000 \times 10^{-3} s^{-1}$ |
| $k_s^+$ | $S_U$ binding on $PD_U$ | $1.000 \times 10^{-3} s^{-1}$ |
| $k_s^-$ | $S_U$ unbinding from $PD_US_U$ | $8.925 \times 10^{-2} s^{-1}$ |
| $k_u^+$ | ubiquitin chain binding | $1.000 \times 10^{-3} s^{-1}$ |
| $k_u^-$ | ubiquitin chain unbinding | $5.625 \times 10^{-2} s^{-1}$ |
| $k_t^+$ | transition from $PD$ to $PD_t$ | $5.400 \times 10^{-1} s^{-1}$ |
| $k_t^-$ | transition from $PD_t$ to $PD$ | $10.125 s^{-1}$ |

**Table S2.** The parameters in Eqs. S2 for the simplified model for USP14-regulated proteasomal degradation demonstrated in Fig. 4A.

| parameter | corresponding process | Value |
| --- | --- | --- |
| $k_{su2}^+$ | $S_{U2}$ binding on $PD$ | Same as Tab. S1 |
| $k_{su2}^-$ | $S_{U2}$ unbinding from $PDS_{U2}$ | Same as Tab. S1 |
| $k_i^+$ | tail insertion | Same as Tab. S1 |
| $k_i^-$ | reverse process of tail insertion | Same as Tab. S1 |
| $k_d^+$ | deubiquitination by USP14 | Same as Tab. S1 |
| $k_d^-$ | reverse process of deubiquitination by USP14 | Same as Tab. S1 |
| $k_h^+$ | substrate translocation and hydrolysis | Same as Tab. S1 |
| $k_h^-$ | reverse process of substrate translocation and hydrolysis | Same as Tab. S1 |
| $k_s^+$ | $S_U$ binding on $PD_U$ | Same as Tab. S1 |
| $k_s^-$ | $S_U$ unbinding from $PD_U S_U$ | Same as Tab. S1 |
| $k_u^+$ | ubiquitin chain binding | Same as Tab. S1 |
| $k_u^-$ | ubiquitin chain unbinding | Same as Tab. S1 |
| $k_t^+$ | transition from $PD$ to $PD_t$ | Same as Tab. S1 |
| $k_t^-$ | transition from $PD_t$ to $PD$ | Same as Tab. S1 |

**Table S3.** The parameters in Eqs. S3 for the reactions of substrate degradation without USP14 in Fig. 4B.

| parameter | corresponding process | Value |
| --- | --- | --- |
| $l_{su2}^+$ | $S_{U2}$ binding | Same as $k_{su2}^+$ in Tab. S1 |
| $l_{su2}^-$ | $S_{U2}$ unbinding | Same as $k_{su2}^-$ in Tab. S1 |
| $l_i^+$ | tail insertion | Same as $k_i^+$ in Tab. S1 |
| $l_i^-$ | reverse process of tail insertion | Same as $k_i^-$ in Tab. S1 |
| $l_d^+$ | deubiquitination by RPN11 | $0.909 \text{ s}^{-1}$ |
| $l_d^-$ | reverse process of deubiquitination by RPN11 | Same as $k_d^-$ in Tab. S1 |
| $l_h^+$ | substrate translocation and hydrolysis | Same as $k_h^+$ in Tab. S1 |
| $l_h^-$ | reverse process of substrate translocation and hydrolysis | Same as $k_h^-$ in Tab. S1 |

### Equations S1 for the simplified kinetic model in Fig. 1C

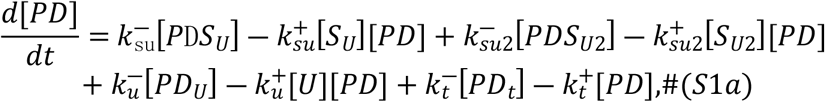

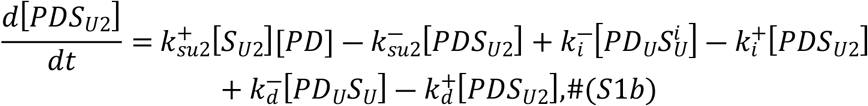

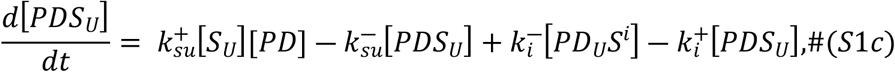

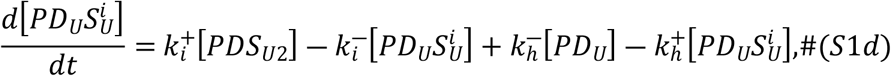

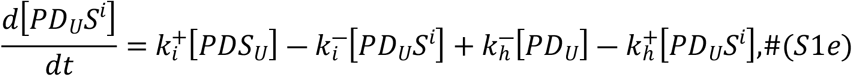

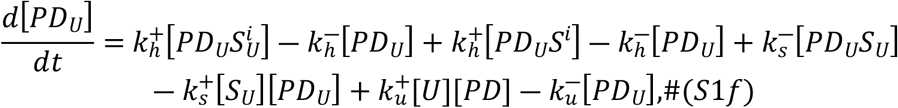

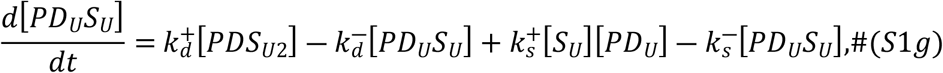

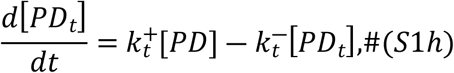

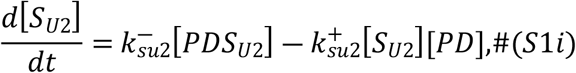

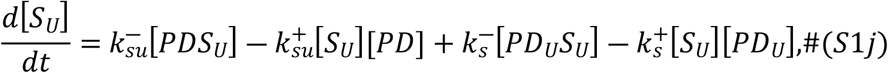

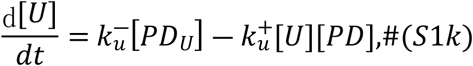

where [.] denotes the concentration of a state variable, and *k*’s are the rate constants. Superscript signs + or – in *k*’s are for forward and reverse rate constants within a reversible reaction. 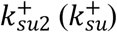 and 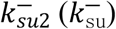 are the rate constants for S_*U*2_ (*S*_U_) binding and unbinding on E_A_-like conformations, while 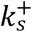 and 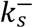 are the rate constants for *S*_*U*_ binding and unbinding on S_D_-like conformations. 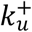 and 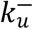 are the rate constants for ubiquitin chain binding and unbinding. Similarly, subscripts i. *d. h. t* correspond to the reactions of substrate insertion, USP14 deubiquitination, substrate hydrolysis, and the transition between *PD* and *PD*_*t*_, respectively. The detailed reactions and rate constants are listed in Tab. S1.

### Equations S2. ODEs for the simplified kinetic model in Fig. 4A

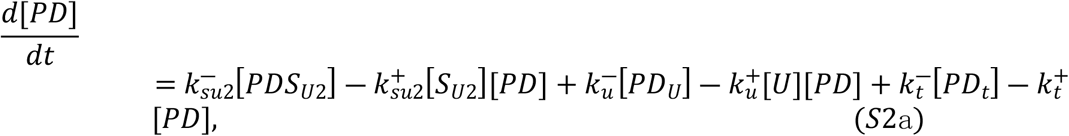

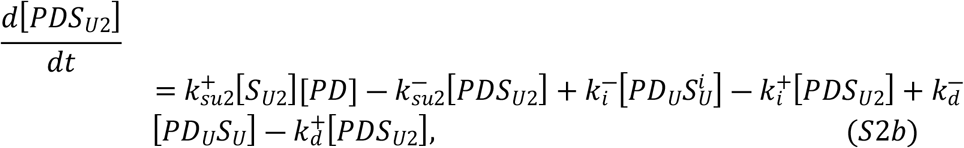

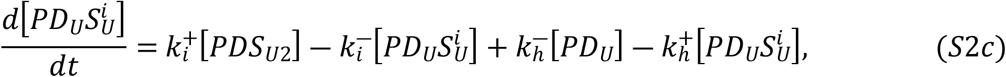

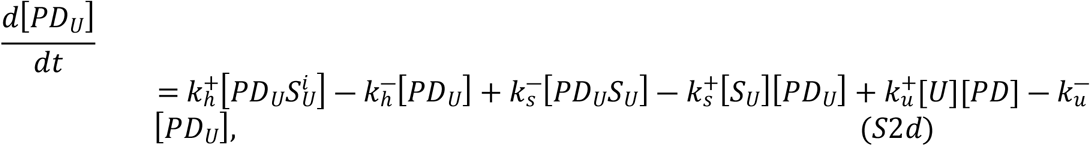

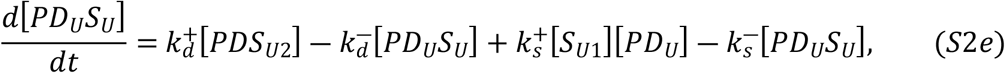

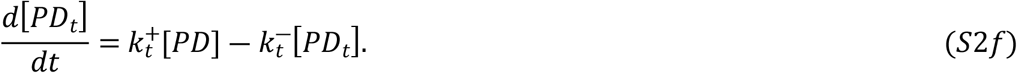

### Equations S3. ODEs for the reactions of substrate degradation without USP14 as shown in Fig. 4B

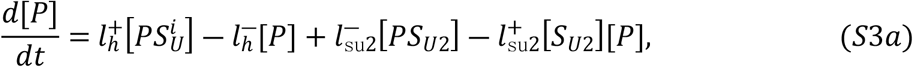

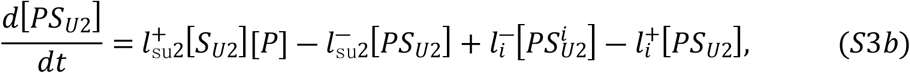

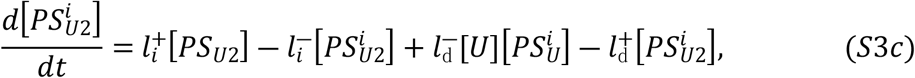

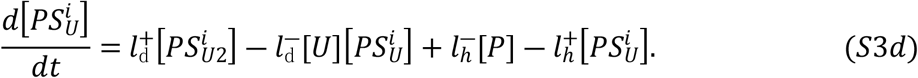

where the rate constants are denoted by *l* to distinguish from the rate constants of the USP14-bound proteasome that are denoted by *k*. Note that *l*_*d*_ denotes the deubiquitination rate of RPN11, which differs from the deubiquitination rate indicated by *k*_*d*_ for USP14.

### Equations S4, ODES for USP14-bound and USP14-free proteasome with two substrates *S* and *T*

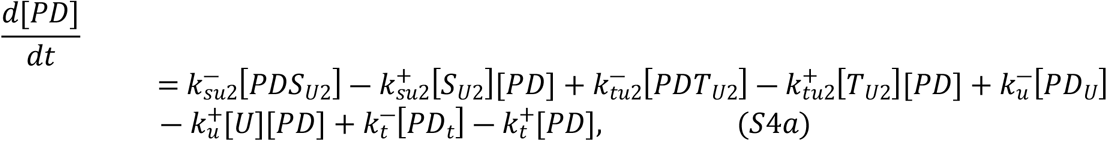

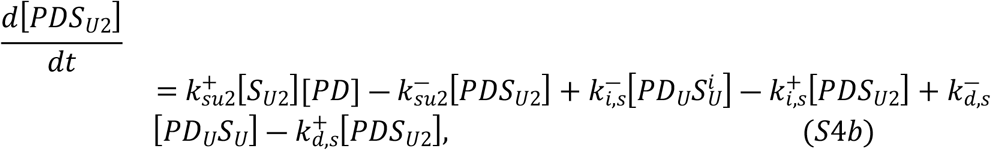

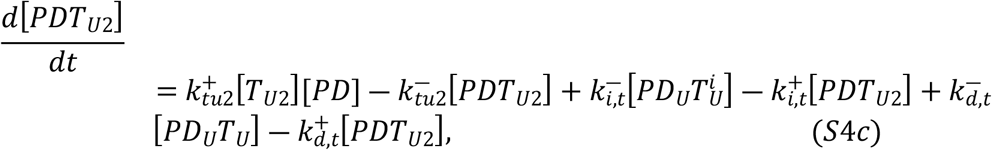

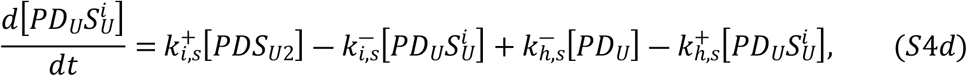

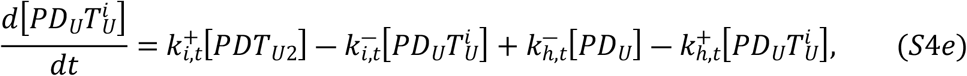

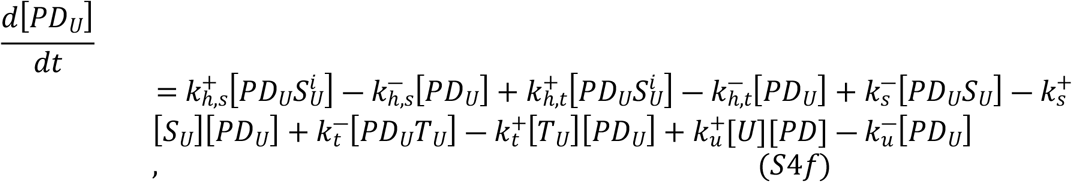

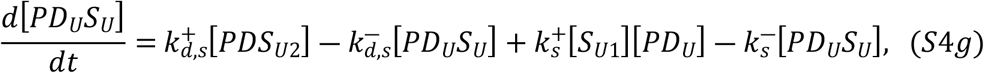

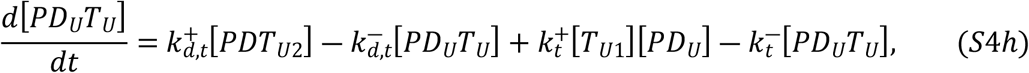

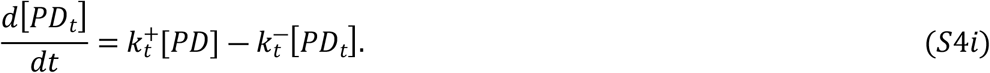

### Equations S5. ODES for USP14-free proteasome and in the presence of substrates *S* and *T*

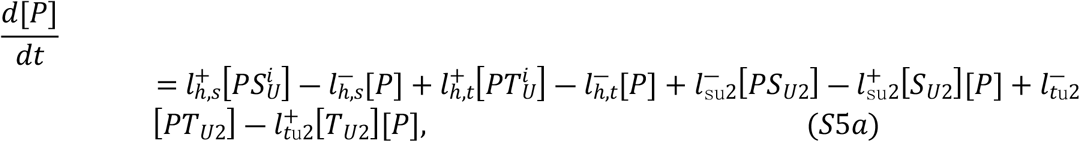

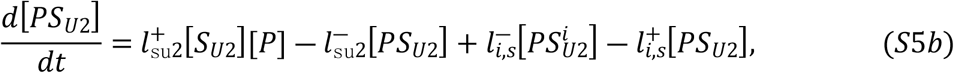

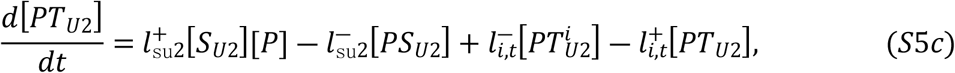

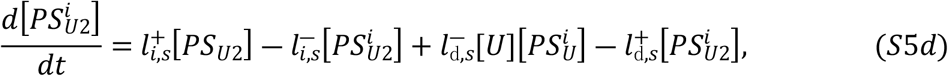

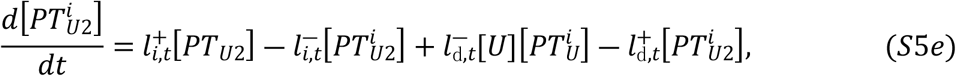

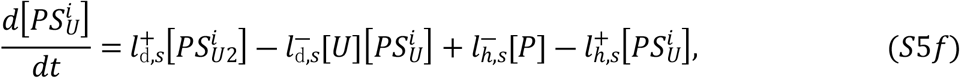

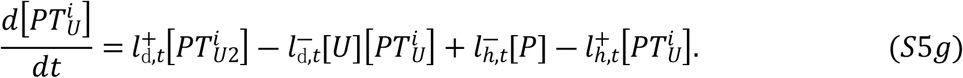

For both Eqs. S4 and S5, subscript *s* and *t* is used to distinguish the corresponding reactions for substrates *S* and *T*, respectively.

### Fig. S1. Parameter sensitivity analysis for the dynamics of USP14-bound proteasome described by Eqs. S1

**Fig. S1.**
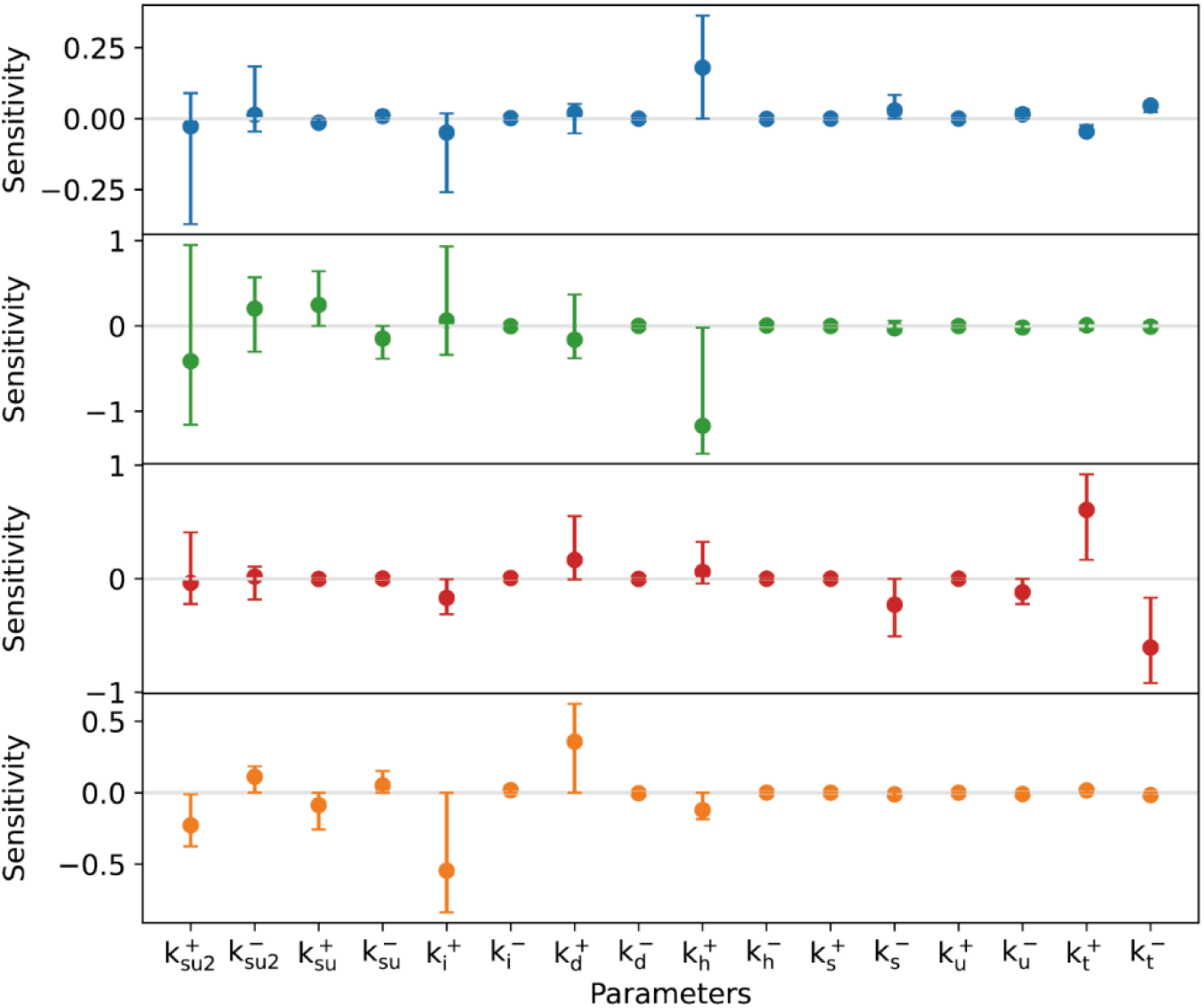
Parameter Sensitivity Analysis of the kinetic model (Eqs. S1) for USP14-bound Proteasome. The sensitivity is defined as *s*(*t*;*p*) ≡ ∂*lnX*(*t*;*p*) ∂*lnp*. From top to bottom: the sensitivities of E_A_-like, E_D_-like, and S_D_-like conformations, as well as the sensitivity of the residual substrate concentration ratio to perturbations in various parameters. The points represent the average sensitivity, with error bars indicating the range of sensitivities. The most sensitive parameters are the substrate binding and unbinding rates 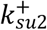 and 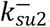, the tail insertion rate 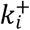, the USP14 deubiquitination rate 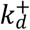, and the substrate hydrolysis rate 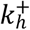.

